# Proteogenomics analysis to identify acquired resistance-specific alterations in melanoma PDXs on MAPKi therapy

**DOI:** 10.1101/2022.02.15.480454

**Authors:** Kanishka Manna, Prashanthi Dharanipragada, Duah Alkam, Nathan L. Avaritt, Charity L. Washam, Michael S. Robeson, Ricky D. Edmondson, Samuel G. Mackintosh, Zhentao Yang, Yan Wang, Shirley H. Lomeli, Gatien Moriceau, Stephanie D. Byrum, Roger S. Lo, Alan J. Tackett

**Author notes:** contributed equally (Kanishka Manna, Prashanthi Dharanipragada, Duah Alkam, Nathan L. Avaritt).

## Abstract

Therapeutic approaches to treat melanoma include small molecule drugs that target activating protein mutations in pro-growth signaling pathways like the MAPK pathway. While beneficial to the approximately 50% of patients with activating *BRAF*^V600^ mutation, mono- and combination therapy with MAPK inhibitors is ultimately associated with acquired resistance. To better characterize the mechanisms of MAPK inhibitor resistance in melanoma, we utilize patient-derived xenografts and apply proteogenomic approaches leveraging genomic, transcriptomic, and proteomic technologies that permit the identification of resistance-specific alterations and therapeutic vulnerabilities. A specific challenge for proteogenomic applications comes at the level of data curation to enable multi-omics data integration. Here, we present a proteogenomic approach that uses custom curated databases to identify unique resistance-specific alternations in melanoma PDX models of acquired MAPK inhibitor resistance. We demonstrate this approach with a *NRAS^Q61L^* melanoma PDX model from which resistant tumors were developed following treatment with a MEK inhibitor. Our multi-omics strategy addresses current challenges in bioinformatics by leveraging development of custom curated proteogenomics databases derived from individual resistant melanoma that evolves following MEK inhibitor treatment and is scalable to comprehensively characterize acquired MAPK inhibitor resistance across patient-specific models and genomic subtypes of melanoma.

## Introduction

Customizing therapeutic strategies based on profiling of tumor-specific alterations has begun to inform clinical practice, though providing individualized treatments based on these findings as a standard-of-care requires additional clinical evidence. Furthermore, co-existence of somatic mutations which drive tumorigenesis and alterations that do not directly control cancer initiation and progression confounds identification of therapeutic vulnerabilities for personalized cancer treatment^1^. Next-generation DNA and RNA sequencing technologies have revolutionized our ability to identify germline and somatic mutations that are associated with cancer, while technological approaches like proteomics allows one to understand whether genomic alterations lead to translation of mutated, variant proteins and/or differentially abundant proteins in cancer cells. The merging of cancer genomics, transcriptomics, and proteomics into proteogenomics applications has the potential to identify novel therapeutic vulnerabilities for the development of the next generation of cancer therapeutics^2,3^.

Cutaneous melanoma is a cancer associated with a high mutational burden, making it a particularly opportune disease for proteogenomics studies. Current therapeutic approaches to treat melanoma include small molecule drugs that target activating protein mutations in pro-growth signaling pathways like the MAPK/ERK pathway and immunotherapies that target cell surface immune checkpoints such as CTLA-4, PD-1, and PD-L1^4,5^. While each type of therapy has particular clinical application and benefit, each is ultimately associated with primary and acquired resistance^6^. Thus, approaches are needed to identify new therapeutic vulnerabilities that can decrease resistance to melanoma therapies and promote a more durable response to treatment, improving survival and quality of life for patients.

The benefit of MAPK inhibitor (MAPKi) therapy against melanoma has been limited to the approximately 50% of patients with activating *BRAF*^V600^ mutation^7,8^. Since the successful development of MEK inhibitor (MEKi) combination to suppress acquired resistance to type I RAF inhibitor (RAFi)^4,9^, progress has stalled to further suppress resistance to the combination of type I RAFi plus MEKi. Similarly, there has been little progress in the development of MAPKi therapy for the other approximately 50% of patients with *BRAF*^V600^ wild-type melanoma, due to contraindication of type I RAFi and to rapid resistance development to MEKi monotherapy in this large subset of patients. Recent studies, including ours, support the notion that a wide-spectrum (*BRAF*^V600MUT^, *NRAS*^MUT^, *NF1*^-/-^, etc.) of melanoma is highly addicted to the MAPK pathway^10^. The lack of efficacy of single-agent MAPKi (e.g., MEKi) therapy belies this exquisite pathway addiction because of an inability to pharmacologically control MAPK pathway reactivation. Thus, effective preclinical strategies that directly tackle MAPK pathway reactivation as well as downstream or parallel phenotypes of acquired resistance warrant clinical development.

To better understand tumor-specific signal dysregulations, the Clinical Proteomic Tumor Analysis Consortium (CPTAC) has conducted mass spectrometry-based proteomic characterization of a growing collection of tumors initially characterized by The Cancer Genome Atlas (TCGA)^11,12^. CPTAC efforts have yet to focus on melanoma, including acquired MAPKi-resistant melanoma. Thus, there is a need to understand resistance evolution in melanoma to inform development of combination therapies. TCGA skin melanoma established the main mutational subgroups (*BRAF, NRAS, NF1*, and triple wild-type), but does not fully account for intra-tumoral heterogeneity, which manifests itself prominently during the evolution of MAPKi resistance^9,13-15^. The co-mutation frequencies of significantly mutated genes in TCGA melanoma are low, suggesting few possible combinatorial targets. During the evolution of MAPKi resistance in *BRAF*^V600MUT^ melanoma, the most frequent and best documented genomic drivers are *RAS* (most frequently *NRAS*) co-mutation and high-magnitude *BRAF*^V600MUT^ amplification^9,14-16^. This level of genomic heterogeneity was not resolved by TCGA melanoma, as only a couple of *BRAF/NRAS* co-mutated melanoma were detected. Furthermore, we have come to appreciate the extensive level of non-mutational melanoma phenotypic plasticity in clinical melanoma even just days and weeks after MAPKi therapy^17^.

It is not practical or feasible to use solely clinical tumor samples to identify and study mechanisms of therapeutic resistance in melanoma, given the challenges of serial clinical biopsies and the lack of targeted therapy for patients with *BRAF*^V600WT^ melanoma. Mounting evidence supports patient-derived xenograft (PDX) models as avatars that faithfully recapitulate the genetic, histologic, and clinical phenotypes of the original tumors^18^. Each PDX model retains genetic variability that impacts gene expression and protein abundance in distinct tumors with acquired resistance to treatment. This genetic variability can be manifested into dysregulation of wild-type protein abundance as well as proteins with amino acid mutations that may serve as novel therapeutic vulnerabilities. We have developed a bank of PDX models that represent a wide range of genomic subtypes in melanoma and chose one model of *NRAS^Q61L^* melanoma to develop this novel proteogenomic approach^10,19^. Furthermore, we have begun to leverage these melanoma PDX models to study mechanisms of acquired MAPKi-resistance^10,19^.

To identify acquired resistance-specific alterations (RSAs) in this *NRAS^Q61L^* melanoma PDX model treated *in vivo* with MAPKi, we applied proteogenomic approaches that leverage genomic, transcriptomic, and proteomic technologies to identify potential therapeutic vulnerabilities. A specific challenge for proteogenomics approaches comes at the level of data curation to enable multi-omics data analysis and integration. Specifically, approaches for the development of uniformly annotated and curated proteogenomic databases are needed to support downstream multi-omics data integration. Here, we present a proteogenomics approach leveraging the development of patient/model-specific curated proteogenomics databases to identify RSAs in melanoma PDX tumors with acquired MAPKi-resistance (**Figure 1A**). Our strategy leverages development of a custom curated proteogenomics database derived from individual acquired-resistant melanoma tumors following MEKi (trametinib) treatment. This approach supports the identification of new targets for future therapeutic development, targeting mechanisms of acquired resistance in distinct genomic subtypes of melanoma.

**Figure 1.**
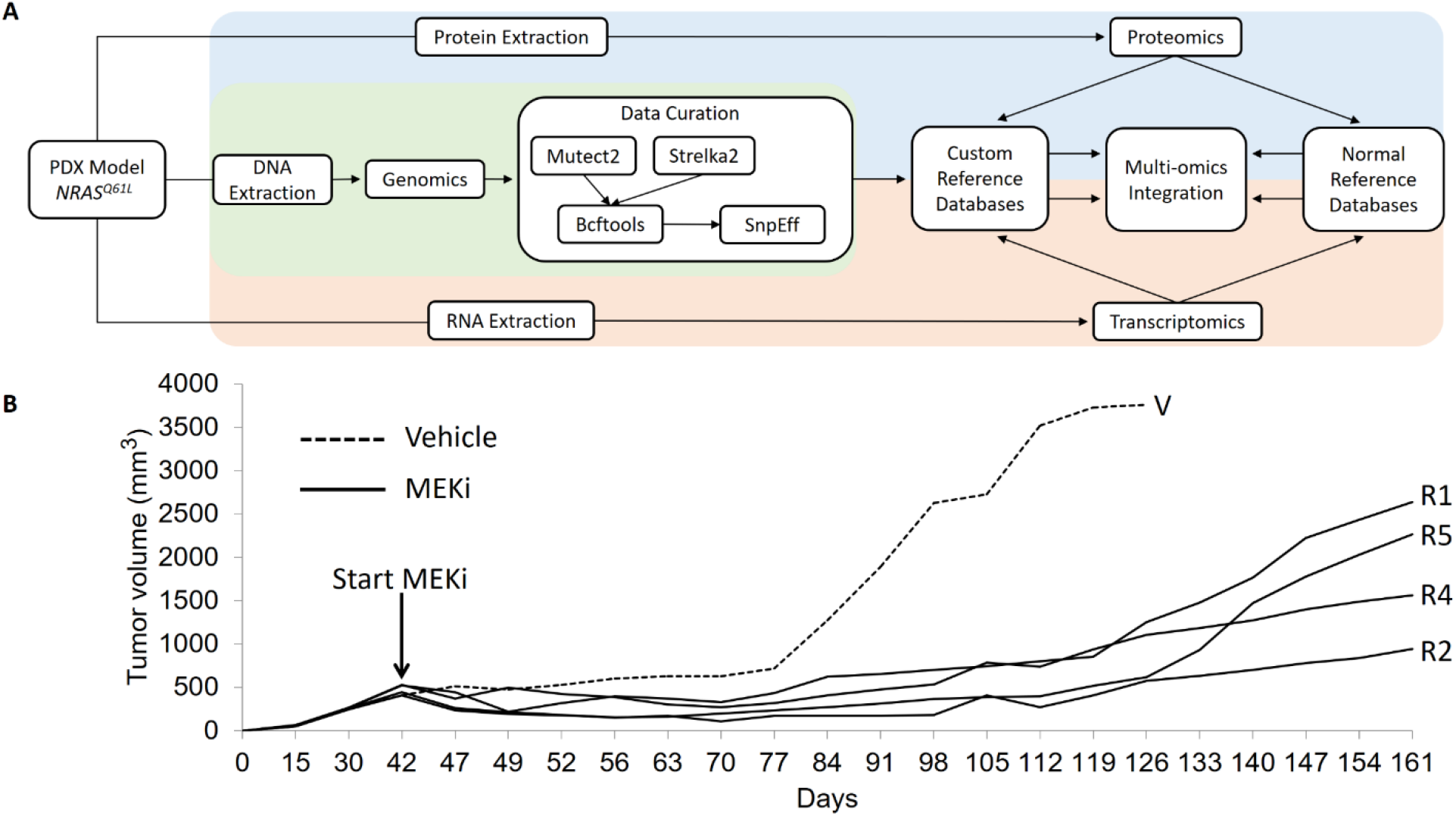
Proteogenomics pipeline for identification of unique resistance-specific alterations and therapeutic vulnerabilities using a PDX model of acquired MEKi resistance. **(A)** Proteogenomics pipeline for genomics (WES), transcriptomics (RNA-seq), proteomics (TMT), data curation, and multi-omics data integration. (**B**) A *NRAS^Q61L^* patient melanoma was established in a NSG mouse model and resulting PDXs were treated with vehicle control or MEKi (trametinib) once tumors reached a volume of approximately 500 mm^3^. At end-point, tumors were prepared for WES, RNA-seq, and proteomics analysis. R: MEKi-acquired resistance; V: vehicle-treated.

## Results and Discussion

### Melanoma NRAS^Q61L^ PDX model

A PDX of an *NRAS^Q61L^* patient melanoma tumor (Mel_PDX2) was established as non-dissociated tumor fragments in NOD scid gamma (NSG) mice and serially passaged without dissociation, in vitro growth cycles, or adjunctive in vivo growth supplements such as Matrigel. To derive the acquired-resistant *NRAS^Q61L^* tumors, we implanted one tumor fragment (passage #2) per NSG mouse. A cohort of tumor-bearing mice with similar tumor volumes were selected for experimentation. When the tumors reached ~500 mm^3^, a mouse was treated with the vehicle-chow, while four mice were treated with trametinib-embedded chow to achieve 5 mg/kg/day dosing (**Figure 1B**). All trametinib-treated tumors displayed tumor regression at this chosen dosage. Tumor tissues for *NRAS^Q61L^* vehicle and four treated samples were harvested at end-points and prepared for whole exome sequencing (WES), bulk RNA-sequencing (RNA-seq), and quantitative tandem mass tag (TMT) proteomics analysis (**Figure 1**).

### PDX tumors with acquired MEKi-resistance show a similar pattern of somatic mutations

Genomic DNA (gDNA) from four acquired-resistant and one vehicle tumor, along with gDNA from a matched normal tissue, were subjected to WES sequencing. Using WES data from the normal patient sample in comparison to each of the PDX tumors, Mutect2 and Strelka2 variant callers were used to assess the somatic mutations in vehicle- and MEKi-treated/acquired-resistant *NRAS^Q61L^* melanoma tumors. The most common SNV class resulted in T>A and C>T nucleotide variants, with SNPs and deletions identified as the most common variant type across tumors (**Figure 2A-E**). Among tumors, missense mutations were identified as the most common variant classification followed by in-frame deletions (**Figure 2A-E**). The frequency of missense mutations and in-frame deletions were consistent across the genes containing the highest levels of mutations (**Figure 2A-E**). Vehicle and resistant tumors shared an overall high level of similarly mutated genes (6,803 genes), but each tumor also had uniquely mutated genes (**Figure 2F**). 250 mutated genes were shared by all four resistant tumors but not the vehicle tumor (**Figure 2F**). The development of unique mutations across resistant tumors relative to vehicle illustrates how genetic divergence occurs as a result of acquired MAPKi-resistance in melanoma. The somatic mutation data reveals similarity, but also uniqueness, across the vehicle and resistant tumors, which supports our rationale for assembly of unique proteogenomics reference databases for transcriptomic and proteomic studies for each resistant tumor compared to vehicle. The creation of model- and tumor-specific proteogenomics databases will more comprehensively enable the rigorous identification of common RSAs and therapeutic vulnerabilities.

**Figure 2.**
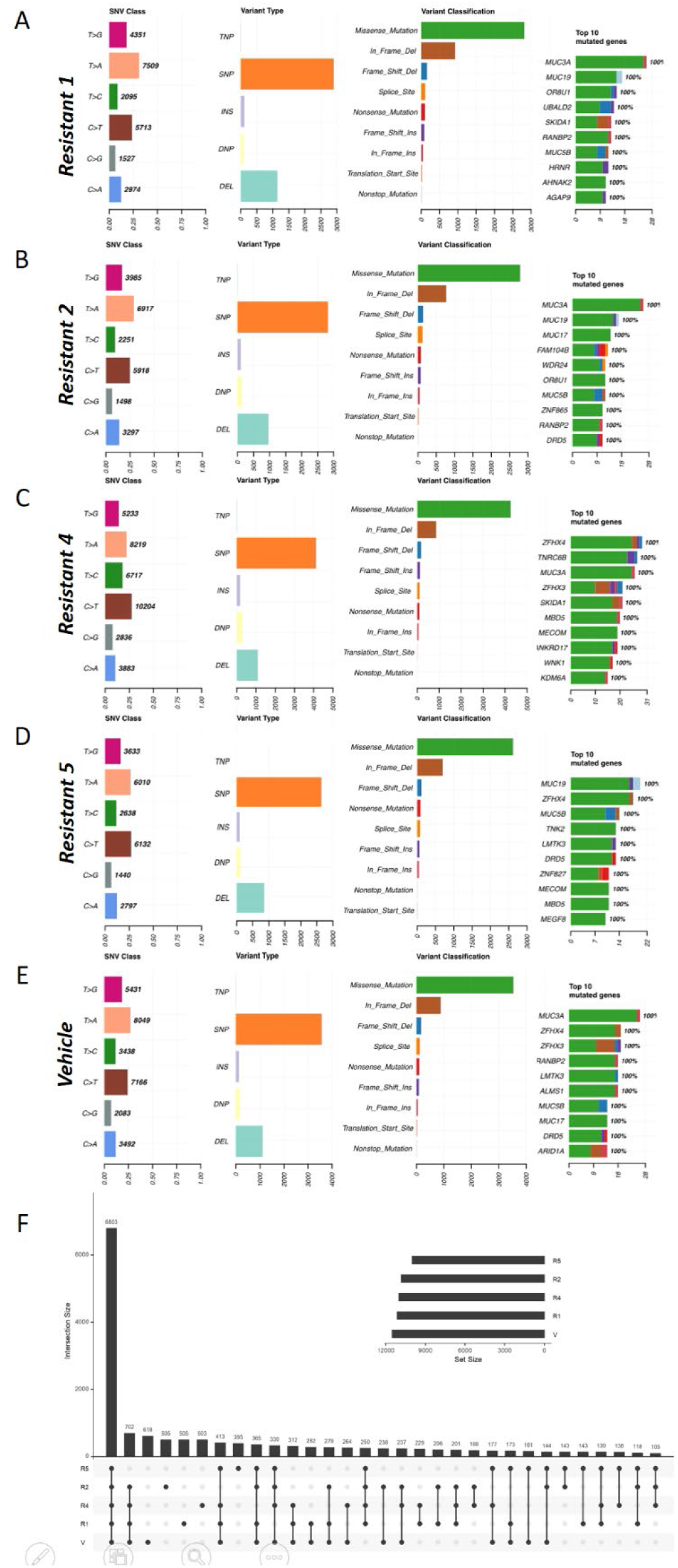
PDX tumors with MEKi-acquired resistance show a similar pattern of somatic mutations. With tumor specific WES data, Mutect2 and Strelka2 variant callers were used to assess somatic mutations in MEKi- **(A-D)** or vehicle-treated **(E)** *NRAS^Q61L^* melanoma. SNV class, variant type, variant classification, and top ten most mutated genes are shown for each tumor. **(F)** UpSet intersection diagram comparing all genes with a detected mutation among MEKi- and vehicle-treated models. R: MEKi-acquired resistance; V: vehicle-treated.

### Correlation of gene and protein data identifies common RSAs

We investigated the expression and abundance levels of genes and proteins using RNA-seq and TMT quantitative proteomics. RNA-seq reads were mapped to the hg38 reference genome (accession: GCA_000001405.15, see methods for details) and the log2 fold change values were calculated in order to identify up-regulated and down-regulated genes between R1 vs V, R2 vs V, R4 vs V, and R5 vs V. Similarly, proteins were identified by searching against the UniprotKB database restricted to *Homo sapiens* (see methods for details). Log2 fold change values were calculated for each resistant tumor verses vehicle, similar to RNA-seq analysis.

The RNA-seq Ensembl IDs were matched to entrez IDs and UniprotIDs using BioMart in order to combine gene expression and protein abundance data. Genes and proteins were then linked by the UniprotID, which was the same identifier in both datasets. RNA-seq data contained 13,210 total genes identified and of these 11,059 genes contained a UniprotID (**Supplemental Table 1**). A total of 7,821 proteins were identified from the protein analysis. Some of these proteins were only identified in the vehicle or the resistant tumors and therefore no fold change is provided (**Supplemental Table 2**). Combining gene and protein molecules using the UniprotID and requiring a fold change value between the resistant tumors compared to vehicle resulted in an integrated data set of 7,229 molecules for a Pearson correlation analysis (**Supplemental Tables 3-6**).

Pearson correlation was performed on the RNA-seq and proteomic log2 fold change values in order to determine consistency between gene expression and protein abundance among the resistant and vehicle-treated tumors. Each data point on the correlation plot in Figure 3 is defined as a detected protein with associated gene. The dotted lines represent a fold change of 2-fold up-regulated (positive) or down-regulated (negative) values. Each of the nine graph subsections (**Figure 3A-D**) indicate molecules that are positively correlated in expression between genes and proteins (right_top and left_bottom), negatively correlated (right_bottom and left_top subsection), up-regulated only in protein abundance (right_middle), up-regulated only in gene expression (middle_top), down-regulated only in protein (left_middle), down-regulated only in gene expression (middle_bottom), or no change in either protein or gene expression (middle_middle). The subsections are defined as left, middle, right along the protein log2 fold change x-axis and top, middle, bottom along the gene log2 fold change y-axis. The proteins and genes showing > 2-fold change when comparing a resistant tumor to vehicle are defined as RSAs and putative therapeutic vulnerabilities. Molecules in the top right subsection (highlighted in red) are up-regulated transcripts and proteins (58 up-regulated proteins with associated up-regulated gene expression for R1 vs V, 50 for R2 vs V, 55 for R4 vs V, and 59 for R5 vs V) (**Figure 3A-D**). Molecules in the bottom left subsections (highlighted in blue) are transcripts and proteins both being down-regulated (89 down-regulated proteins with associated down-regulated gene expression for R1 vs V, 64 for R2 vs V, 28 for R4 vs V, and 87 for R5 vs V) (**Figure 3A-D**). Molecules in the middle right subsections (highlighted in purple) are up-regulated protein abundance with no significant change in gene expression (183 up-regulated proteins for R1 vs V, 153 for R2 vs V, 84 for R4 vs V, and 153 for R5 vs V) (**Figure 3A-D**). Molecules in the middle left subsections (highlighted in yellow) are down-regulated protein with no significant change in gene expression (108 down-regulated proteins for R1 vs V, 76 for R2 vs V, 32 for R4 vs V, and 129 for R5 vs V) (**Figure 3A-D**). The latter two classes of RSAs suggest post-transcriptional proteomic alterations. Across all comparisons of resistant to vehicle tumors, few examples of up-regulated genes with down-regulated proteins (left_top subsections) and down-regulated genes with up-regulated proteins (right_bottom subsections) were identified (**Figure 3A-D**). Positive correlation of gene and protein levels provides a measure of validation for a RSA that could be a therapeutic vulnerability, while those molecules with a negative correlation between gene and protein are also putative therapeutic vulnerabilities due to the well-recognized discordance of gene and protein levels^11,12,20^.

**Figure 3.**
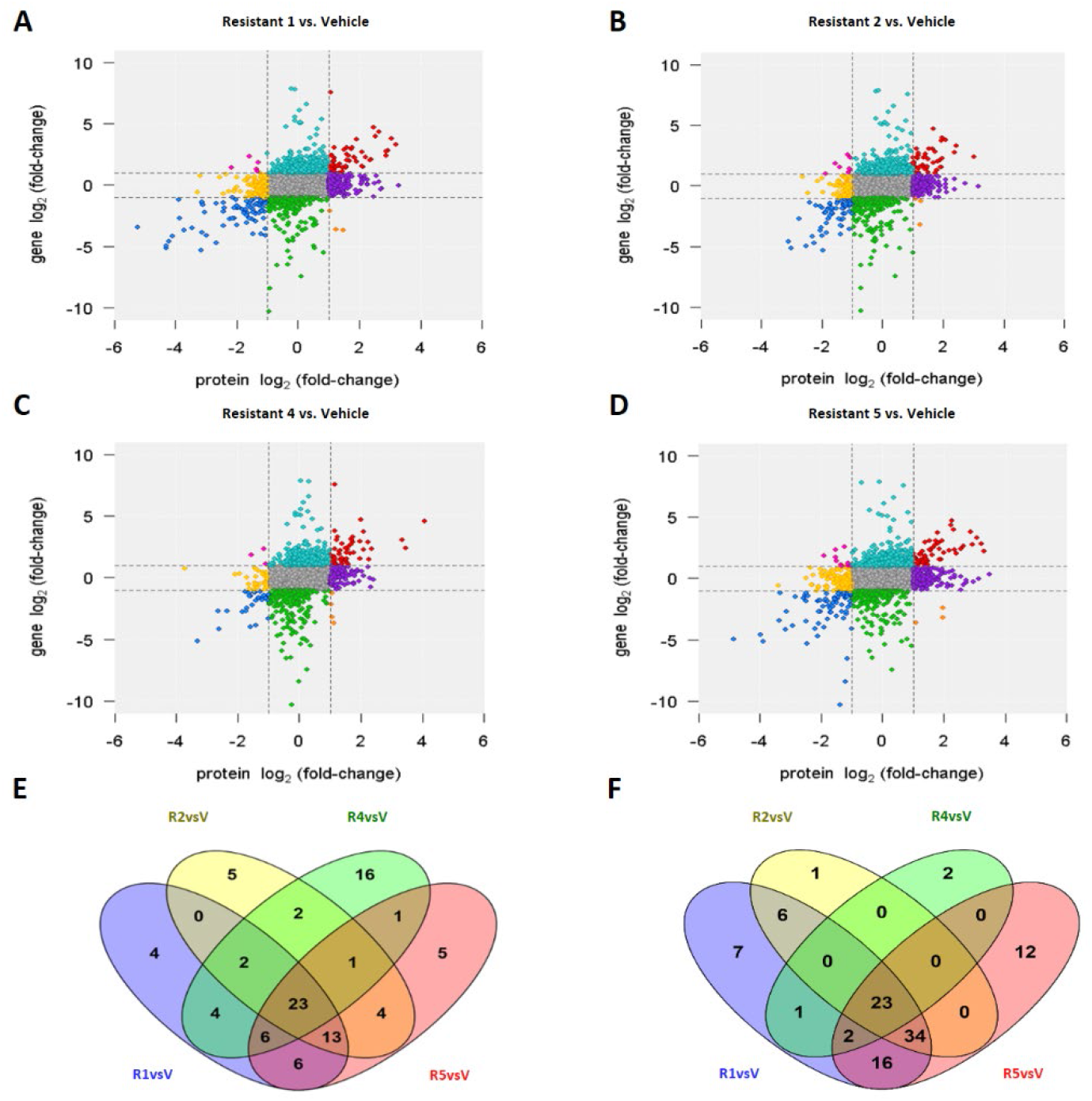
Pearson correlation of transcriptome and proteome data identifies common resistance-specific alterations and vulnerabilities across PDX models of MEKi-acquired resistance. **(A-D)** The log2 fold change values for each resistant tumor compared to vehicle is correlated for RNA-seq and proteomics data. Molecules with >2-fold change are indicated by dashed lines. **(E,F)** Venn diagrams showing molecules up-regulated **(E)** or down-regulated **(F)** at both the transcript and protein level across resistant tumors relative to vehicle control. R: MEKi-acquired resistance; V: vehicle-treated.

To identify common RSAs shared across all resistant tumors relative to vehicle control, we compared molecules that were up-regulated at both the gene and protein levels (identified in the top right subsection and highlighted red) and down-regulated at both gene and protein levels (identified in the bottom left subsection and highlighted blue). Out of the lists of molecules in each resistant tumor compared to vehicle a total of 23 molecules were consistently up-regulated and down-regulated. (**Figure 3E, F**). The 23 shared across all resistant tumors for up-regulated transcripts and proteins were *MATN2, HNMT, HPGD, SYNPO, SELENBP1, DCLK1, HMCN1, EDNRB, TPCN2, MET, SOX6, ABCC2, P2RX7, FAM129A, BCAN, PRELP, CPS1, PPFIBP2, SULT1A4, TTF2, COBLL1, CRYAB*, and *AHNAK*. The 23 shared across all resistant tumors for down-regulated transcripts and proteins were *CDKN2B, COL4A1, COL4A2, CYB561, DUSP4, ISG20, JAG1, P4HTM, RFTN1, SERPINE1, TMEM102, TMEM163, TUSC3, USP53, VGF, ASNS, ATP1B1, CADPS, FCGRT, MAGEA10, NELL1, OGDHL*, and *TUBA8*. At the level of concordant gene and protein levels, all four resistant tumors displayed a high level of convergent evolution, with R4 resistant tumor being most divergent. The common and unique RSAs identified in **Figure 3** support the need for using a custom proteogenomics database for each specific tumor.

### Custom proteogenomics reference databases enable mutanome-specific identification of RSAs and pinpoint a driver alteration

To create a custom proteogenomics database for each resistant tumor model, the intersection of variants identified using Mutect2 and Strelka2 from WES data was used to build custom transcriptomic and proteomic databases to identify mutated genes and proteins not present in the standard human reference databases (**Figure 1A**). The variant calling files were annotated using SnpEFF; and the variant annotation was used to create uniformly curated, custom genome and protein fasta files that included nucleotide and protein amino acid changes. The uniform annotation within the database enables more straightforward and rigorous downstream multi-omics analyses. The majority of variants identified in the WES data were missense mutations, which result in amino acid changes in proteins that may impact function and stability. The RNA-seq and proteomic data sets were searched using the custom proteogenomic databases for each tumor (**Supplemental Tables 7 and 8**). We identified a total of 190 variant proteins that had associated gene transcription information across all tumor models. Of those detected, 47 variant proteins showed an absolute 1.5-fold difference comparing all resistant models to vehicle. The z-score of the log2 normalized gene expression values and protein abundances are displayed in **Figure 4A**. The RSAs in **Figure 4A** are sorted by highest to lowest protein level in the resistant tumors compared to vehicle. The protein amino acid mutation is also indicated. By using the custom proteogenomics database, we were able to identify the variant RSAs found in **Figure 4** which were not identified using the standard database searches utilized in **Figure 3**, demonstrating the value of this novel proteogenomic approach.

**Figure 4.**
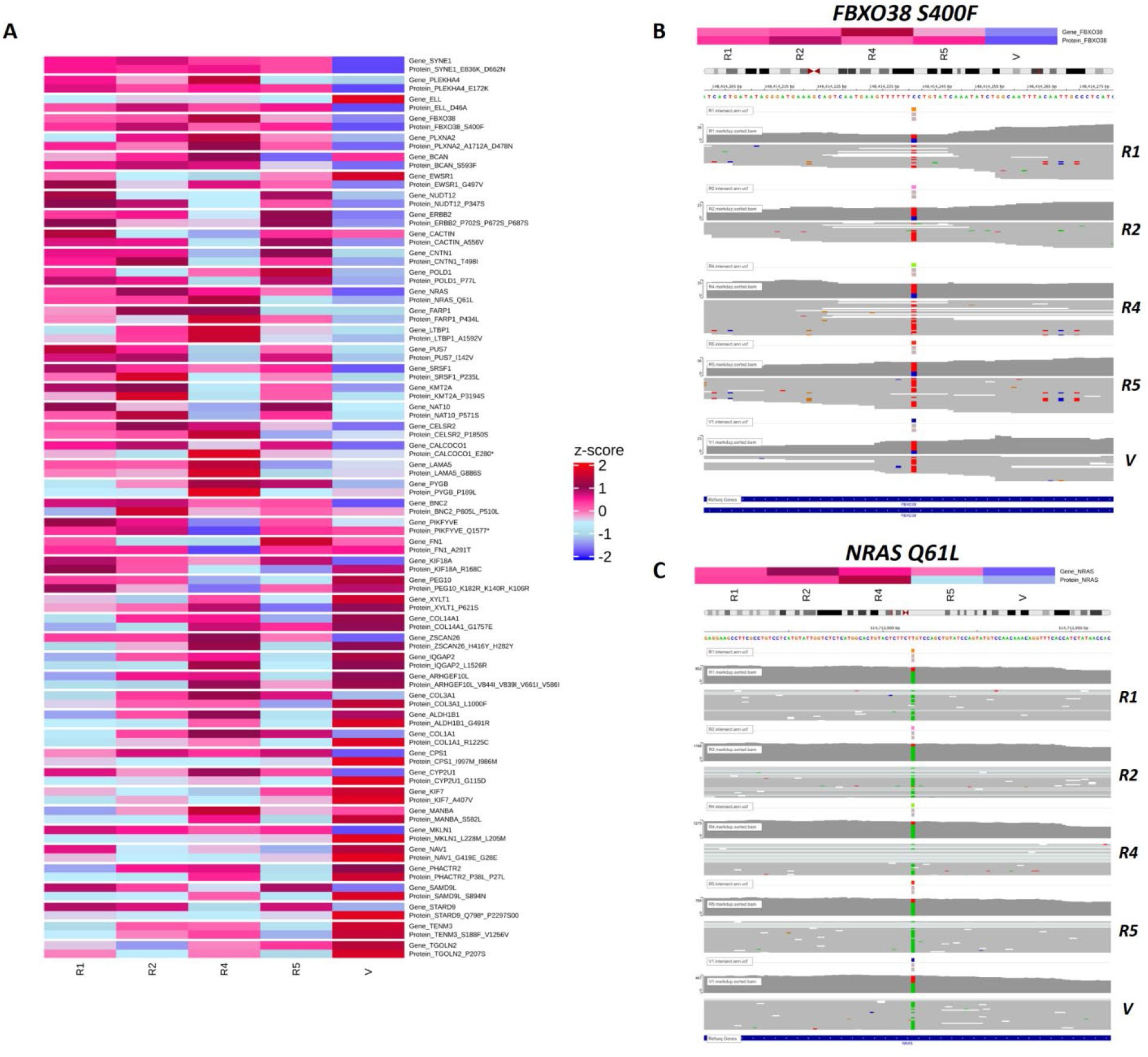
Custom proteogenomic reference databases enable identification of unique resistance-specific alterations and vulnerabilities across PDX models of MEKi-acquired resistance. **(A)** RNA-seq and proteomics data were searched against the custom proteogenomics databases assembled from WES data. The log2 normalized values for the differentially regulated proteins and associated gene transcripts are shown (>1.5 fold-change). The amino acid change is shown for each protein. **(B,C)** Expanded view of RNA-seq data for FBXO38 **(B)** and NRAS **(C)**. WES variant calling files and RNA-seq coverage tracks are shown for each vehicle and resistant tumor. R: MEKi-acquired resistance; V: vehicle-treated.

As examples of putative vulnerabilities uniquely identified with our custom proteogenomics database strategy, we highlight FBXO38 and NRAS (**Figure 4B, C**). The F-box protein FBXO38 is a member of the SKP1-CUL1-F-box protein (SCF) family of E3 ubiquitin ligases, which have been implicated in cancer-associated drug resistance^21^. Using our custom proteogenomics databases, FBXO38 was shown to have elevated gene and protein levels across all resistant tumors relative to vehicle (**Figure 4B**). WES data showed FBXO38 having a genomic C>T missense mutation leading to a Ser400Phe protein amino acid mutation. This somatic mutation, RNA-seq read coverage, and RNA-seq individual reads mapping to chromosome 5 position 148,414,241 are illustrated in **Figure 4B**. The RNA-seq reads at FBXO38^S400F^ contain the following percentages of C and T: R1- 50% C, 50% T; R2-31% C, 69% T; R4- 33% C, 67% T; R5- 33% C 67% T; V1- 25% C, 75% T. Similarly, through the use of the custom proteogenomic database, NRAS was shown to have elevated gene and protein levels across all MEKi resistant NRAS^Q61L^ tumors relative to vehicle (**Figure 4C**). The NRAS somatic mutation, RNA-seq read coverage, and RNA-seq reads mapping to chromosome 1 position 114,713,908 is displayed for each sample (**Figure 4C**). The RNA-seq reads at NRAS^Q61L^ the following percentages of T and A: R1- 76% A, 24% T; R2-87% A, 12% T; R4- 84% A, 16% T; R5- 79% A 21% T; V1- 60% A, 40% T. FBXO38 and NRAS proteins were identified as differentially expressed only when searching against the custom proteogenomics database (**Figure 4A**). Importantly, we have shown that *NRAS*^Q61L^ up-regulation (at both the gene and protein level) causes acquired MEKi resistance, driven in some resistant tumors by gene amplification^10^.

In summary, we present a proteogenomics approach that can be applied to PDX models of acquired MAPKi resistance to enable the identification of unique RSAs and putative therapeutic vulnerabilities. Here, we demonstrate the utility of this multi-omics pipeline with a *NRAS^Q61L^* melanoma PDX model with resistant tumors developed following treatment with MEKi. Our results support the need to create individualized, custom proteogenomics databases for each resistant tumor. Furthermore, our data highlights the utility of using both standard databases and custom proteogenomics databases to identify RSAs and therapeutic vulnerabilities across PDX tumors. In the future, we will extend this multi-omics platform across a variety of genetically diverse melanoma PDX models of resistance to MEKi and other MAPKi therapies, which will support the identification of new RSAs for future therapeutic development targeting mechanisms of acquired resistance in genetically distinct subtypes of melanoma. Additionally, future developments will feature user-friendly, interactive HTML outputs for multi-omics data analysis that will support proteogenomics studies beyond those in PDX models of therapy resistance.

## Materials and Methods

### Mice

NSG (NOD scid gamma) mice were obtained from the Radiation Oncology breeding colony at UCLA (Los Angeles, CA). Sex-matched mice were used at 4-6 weeks of age. All animal experiments were conducted according to the guidelines approved by the UCLA Animal Research Committee.

### PDX, Treatment and Tissue Collection

To develop PDX models, tumor fragments derived from NRASQ61L metastatic melanoma (Mel_PDX2), with approval by the local Institutional Review Boards, were transplanted subcutaneously in sex-matched NSG mice (4-6 week old). One tumor fragment was implanted in each mouse. Tumors were measured with a caliper every 2 days, and tumor volumes were calculated using the formula (length x width2)/2. Tumors with tumor volumes around 500 mm^3^ were randomly assigned into experimental groups. Special mice diet (for NSG) was generated by incorporating trametinib at 5 mg/kg to facilitate daily drug dosing at 5 mg/kg/day and to reduce animal stress (Test Diet Richmond, IN, USA). At time of collection (day 126 and 161 for vehicle and resistant tumors respectively), tumors were excised from mice and subjected to WES, RNA-seq and proteomics analysis.

### Next-generation sequencing for WES and RNAseq

Genomic DNA (gDNA) and total RNA were extracted from snap frozen patient tumor and patient-matched normal tissue, and frozen PDX tumors preserved in RNALater using the QIAGEN AllPrep DNA/RNA Mini Kit and the Ambion mirVana miRNA Isolation Kit. All gDNA were quantified using a NanoDrop Spectrophotometer (Thermo Fisher Scientific), then gDNA size and quality were analyzed using agarose gel electrophoresis to confirm for the presence of an intact high molecular weight band. gDNA libraries were constructed using the Roche NimbleGen SeqCap EZ Human Exome Library Kit v2.0, which captures 64 Mb of the human genome by 2.1 million oligonucleotide probes for coverage of more than 20,000 genes in the human genome (Roche NimbleGen, Inc., Madison, WI). Briefly, after fragmentation of gDNA, the libraries were constructed by end repairing and A-tailing the fragmented DNAs, ligation of adapters, and PCR amplification. After library construction, libraries were quantified for equal molar pooling and paired-end sequenced with a read length of 2×150 bp on the Illumina NovaSeq 6000 S4 platform. All total RNA was quantified using a Nanodrop, then RNA size and quality were measured using Agilent 2100 Bioanalyzer (Agilent Technologies). RNA libraries were constructed using the NuGen Universal Plus mRNA-Seq Kit. Briefly, after fragmentation of total RNA and double-stranded cDNA generation using a mixture of random and oligo(dT) primers, the RNA libraries were constructed by end repairing, adapter ligation, strand selection, and PCR amplification. After RNA library construction, libraries were quantified for equal molar pooling and paired-end sequenced with a read length of 2×50 bp on the Illumina NovaSeq 6000 S2 platform.

### WES analysis and creation of custom-curated proteogenomics databases

Fastq files were trimmed using trimmomatic and reads with Q>30 were kept. Reads were aligned to the hg38 reference genome (accession: GCA_000001405.15) using bowtie2. Somatic mutations were identified using Mutect2^22^ and Strelka2^23^ variant callers for V compared to normal, R1 vs normal, R2 vs normal, R4 vs normal, and R5 vs normal with hg38 as reference. The intersection of Mutect2 and Strelka2 variant calling algorithms was used to identify the variants within each tumor, which were used for further downstream analyses. Additionally, the intersection of variants between resistant tumors was identified to find a common set of variants in the resistance group. The variants were filtered to retain only passing ones. The resulting variants were used to build uniformly annotated, custom hg38 reference genomes using Bcftools (version 1.9)^24^. These custom hg38 genomes were used as references for gene and protein analyses. Additionally, the variant files were uniformly annotated and the resulting protein sequences of called variants were obtained using SnpEff (version 5.0e)^25^. Variant calls were visualized using the Circlize package (version 0.4.13)^26^. Summary of variant calls were visualized using the Maftools package (version 2.2.10)^27^. Somatic mutation calls were manually reviewed in IGV.

### RNA-seq analysis

RNA-seq reads were quality-checked, trimmed, and aligned to the hg38 reference genome (accession: GCA_000001405.15) and our custom hg38 genomes using the Nextflow RNAseq pipeline, nf-core/rnaseq (version 3.4) available at DOI 10.5281/zenodo.1400710. The resulting gene counts were transformed to log2 counts per million (CPM)^28^. Lowly expressed genes were filtered out and libraries were normalized by trimmed mean of M-values^29^. Log2 fold change values were calculated for each resistant tumor compared to vehicle. Genes with an absolute fold change > 2 were considered significant (**Supplemental Tables 1 and 7**).

### Proteomic Analysis

Protein (100 μg) of from each tumor sample was reduced, alkylated, and purified by chloroform/methanol extraction prior to digestion with sequencing grade trypsin^30^. The resulting peptides were labeled using a tandem mass tag (TMT) isobaric label reagent set (Thermo). Labeled peptides were separated into 46 fractions on a 100 x 1.0 mm Acquity BEH C18 column (Waters) using an UltiMate 3000 UHPLC system (Thermo) with a 50 min gradient from 99:1 to 60:40 buffer A:B ratio under basic (pH 10) conditions, and then consolidated into 24 super-fractions. Buffer A contains 0.5 % acetonitrile and 10 mM ammonium hydroxide. Buffer B contains 10 mM ammonium hydroxide in acetonitrile. Each super-fraction was then further separated by reverse phase XSelect CSH C18 2.5 um resin (Waters) on an in-line 150 x 0.075 mm column using an UltiMate 3000 RSLCnano system (Thermo). Peptides were eluted using a 75 min gradient from 98:2 to 60:40 buffer A:B ratio. Here, buffer A contains 0.1% formic acid and 0.5% acetonitrile and buffer B contains 0.1% formic acid and 99.9% acetonitrile. Eluted peptides were ionized by electrospray (2.4 kV) followed by mass spectrometric analysis on an Orbitrap Eclipse mass spectrometer (Thermo) using multi-notch MS3 parameters. MS data were acquired using the FTMS analyzer in top-speed profile mode at a resolution of 120,000 over a range of 375 to 1500 m/z. Following CID activation with normalized collision energy of 31.0, MS/MS data were acquired using the ion trap analyzer in centroid mode and normal mass range. Using synchronous precursor selection, up to 10 MS/MS precursors were selected for HCD activation with normalized collision energy of 55.0, followed by acquisition of MS3 reporter ion data using the FTMS analyzer in profile mode at a resolution of 50,000 over a range of 100-500 m/z. Proteins were identified by searching either the UniprotKB *Homo sapiens* database or our custom proteogenomics database using MaxQuant (Max Planck Institute, version 2.0.1). Proteins were normalized using the MS3 reporter ion intensities by Cyclic Loess from the limma package and the log2 fold change values were calculated (**Supplemental Tables 2 and 8**)^31^.

## Author Contributions and Notes

R.S.L, G.M., P.D., A.J.T., and S.D.B designed research, K.M., D.A., N.L.A, C.L.W, M.R., R.D.E, S.G.M., S.H.L., Z.Y., Y.W., P.D., G.M., and S.D.B performed research and analyzed data; and N.L.A, R.S.L., A.J.T., and S.D.B. wrote the paper. The authors declare no conflict of interest except R.S.L., who receives research funding from Merck, Pfizer, and BMS.

## Acknowledgments

This research was supported by grants from the National Institutes of Health: P20GM121293 (A.J.T), R24GM137786 (A.J.T); R01CA236209 (A.J.T.); R01CA176111 (R.S.L.); R21CA215910 (R.S.L.); R21CA255837 (R.S.L.); 1P01CA168585 (R.S.L.); V Foundational for Cancer Research Translational Award (R.S.L.); the Ressler Family Foundation (R.S.L.).

